# Thin Microfluidic Chips with Active Valves

**DOI:** 10.1101/2023.06.09.544232

**Authors:** Ekta Prajapati, Pravin Shankar Giri, Subha Narayan Rath, Shishir Kumar

**Affiliations:** Indian Institute of Technology, Hyderabad, 502285, India

## Abstract

We report the fabrication of very thin microfluidic active and passive devices on rigid and flexible substrates for sample-space-restricted applications. Thin glass coverslips are commonly used substrates, but these being fragile often crack during experiments, leading to device failure. Here, we used PET as a flexible substrate to fabricate robust thin devices. We proposed a simpler process for PET-PDMS bonding without any silane, adhesive, and/or plasma treatment. We presented the compatibility of the thin devices with a digital in-line holographic microscope (DIHM) as a use case. The substitution of the conventional microscope with DIHM in microfluidic large-scale integrated systems renders simplicity, cost-effectiveness, portability, and miniaturization of the overall system. It also enables a customized and parallel multisite optical observation for a complex microfluidic circuit chip. These chips comprise various microfluidic components made of active microvalves, particularly Quake valves. We also successfully demonstrated the function of microvalves fabricated with our method to regulate the fluidic flow. Thus, are suited to making sophisticated microfluidic circuit chips to fit a variety of applications like organ-on-chip, cell culture, wearable biosensors, pressure sensors, etc.

## Introduction

Microfluidics large-scale integrated (mLSI) systems have impressed many life science researchers with their capabilities to perform the assays with high precision, shorter turn-around times, increased efficiency in data collection, low sample and reagent consumption, etc [1,2]. The optical observations of these mLSI chips are conventionally performed using benchtop light microscopes. The typically used thick microfluidic circuit chips are not always compatible for high-resolution imaging, especially, with high numerical aperture (NA) objective lenses. Hence, the need arises for the development and use of thin microfluidic chips. Along a similar line, thin microfluidic devices are also needed for applications like stimulated Raman scattering microscopy, confocal microscopy, IR spectroscopy, optical and magnetic tweezers, etc [3].

The requirement for thin microfluidic chips also arises from replacing the traditional bulky and expensive optical microscopes with their lensless counterparts, particularly Muscope [4]. Muscope is a miniature on-chip digital in-line holographic microscope (DIHM) [5,6], reported by us - has a form factor of about 4 mm and can be easily integrated within larger systems. Its employment with mLSI systems renders multiple advantages such as improved simplicity, cost-effectiveness, portability, and miniaturization of the overall system. It comprises a microLED display as a compact light source and an image sensor, sandwiching the sample under observation. This simple assembly can be used at different locations of an mLSI circuit chip, allowing a multisite parallel observation, aiding to achieve lab-on-a-chip targets. Another attractive feature of Muscope is the decoupling of the resolution and field-of-view (FOV). Thereby allowing the source and sensor specifications of a Muscope to be customized for different locations on mLSI chips demanding a particular resolution and FOV. These distinguishing features of Muscope make it an apt choice for optical observations in mLSI, like real-time imaging, time-lapse imaging, etc.

When employing Muscope with mLSI, the chip form factor impacts the quality of the captured holograms due to reduced signal-to-noise ratio (SNR). Consequently, deteriorates the resolution of the reconstructed image. Thus, in this work, we focus on the fabrication of thin active and passive microfluidic devices [7]. The thin glass coverslip is a common substrate choice to fabricate thin microfluidic-based devices. Glass coverslips, being fragile, often crack while performing experiments. Thus, demand extra attention and careful skilled manual handling. To resolve this, the flexible substrates can be integrated with microfluidic devices [8,9]. In this work, we chose the inexpensive, optically transparent, and easily available, Polyethylene Terephthalate (PET) [10] as the flexible substrate for our thin microfluidic devices. PET offers biocompatibility, uniformity, thermal and chemical resistance, thus, is suitable for applications like wearable flexible patches for biofluid sampling, manipulation, and bio-sensing [11,12]. Moreover, the biocompatibility of PET can be governed by surface modification treatments, opening opportunities for a plethora of applications like organ-on-chip [13], drug discovery [14], cell culture [15,16], etc.

Many of these biological assays are carried out with mLSI circuit chips comprising different microfluidic components like micropumps [17], micromixers [18,19], multiplexers [20,21], and many more, on a single chip. The micromechanical valves function as the basic building block of an mLSI circuits. With decades of development, many microvalve designs have been reported [22]. Although none of the valves are truly “on-chip”. For the large-scale integration, the pneumatically-controlled elastomeric Quake valve [23] remains the best suited. It provides scalability, modularity, automation, and parallel processing for high-throughput biology. A normally-open Quake valve comprises control and flow microchannels, overlapping each other orthogonally to form a valve. The pressurized fluidic flow in the control channel (CC) occludes the sample flow in the flow channel (FC). It can have 2-layer [24] and 3-layer [25] architecture, based on the number of existing polymer layers. The 3-layer design tunes the valve action more efficiently and is more reliable to form complex mLSI chips over its 2-layer counterpart. Due to the merits it offers, we opted for the fabrication of the thin 3-layer microfluidic active device in the present work.

In this work, we demonstrated the fabrication process of thin 1-layer and 3-layer Quake valve-based microfluidic circuits with the thin glass coverslip (#1.5) and flexible PET sheet substrate. To achieve this, we also reported a simpler recipe to bond PDMS and PET without demanding any chemicals, chemical adhesives, or plasma system. We affirmed the device handling and sample flow regulation in our devices with an indigenous microfluidic controller system (inspired by Stephen Quake’s setup). Since many researchers still use 2-layer valve-based microfluidic circuits, we also proposed a simple procedure to fabricate their thinner version. We also presented the comparison between the compatibility of thick and thin microfluidic devices with Muscope. Here, we also extended the Muscope’s capability to function as a Phase Contrast Microscope (PCM) with absolutely no hardware overhead in its experimental setup. In fact, the amplitude and phase information of the sample retrieved by holographic reconstruction of images were utilized to result in phase-contrasted images.

## Experimental Details

### 1. Thin microfluidics device fabrication

We show the fabrication processes of thin microfluidic devices with single and three PDMS layers. The mold for each PDMS layer was prepared on separate silicon wafers using standard photolithography. A photomask was written (Figure 1 (a-d)) and used to pattern SU8 2015 (13-15 *µ*m thick) on the silicon substrate (Figure 1(e-f)). Fluoropel 800 was spin-coated on the molds to form a hydrophobic and oleophobic coating [Figure 1(g)]. These molds can be used for replica molding for many months, ensuring an easy release of even very thin layers of structured PDMS. The mold preparation is detailed in the supplementary information. All the devices were prepared with PDMS, mixed in the ratio of 1:5 (crosslinker ratio to PDMS) unless stated otherwise. The commonly used ratio of 1:10 works well too.

**Figure 1:**
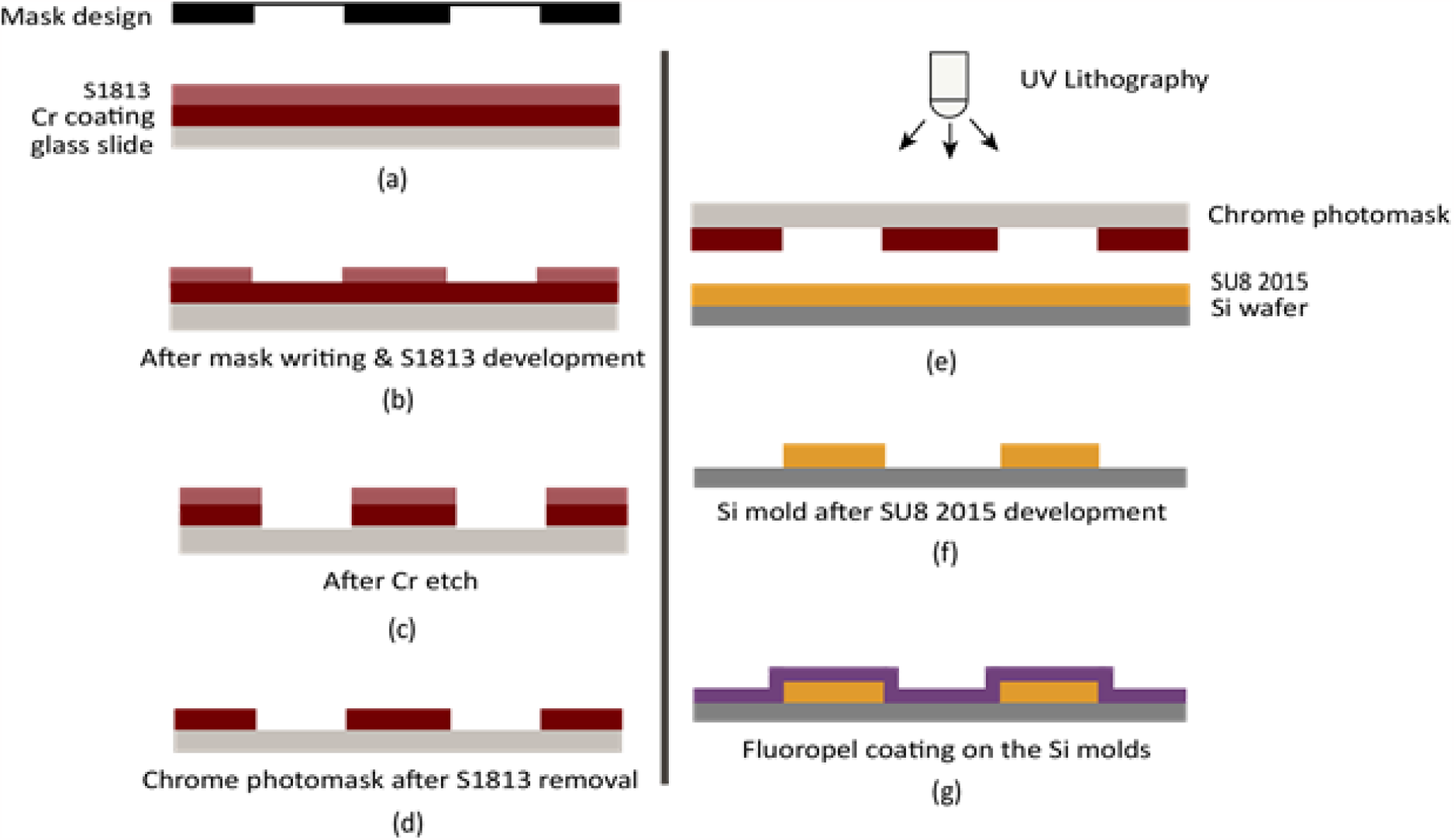
The process flow for chrome mask [(a)-(d)] and silicon mold preparations [(e)-(g)].

#### a) Fabrication of thin 1-layer device

PDMS (1:5) was degassed and spin-coated on the silicon molds. It was cured at 80C for 1.5 hours. The thin structured PDMS layer was carefully peeled off the molds without tearing it. The inlet-outlet ports were pierced with the puncher.

### Glass coverslip as substrate

This structured PDMS was bonded with the glass coverslip (#1.5) using the oxygen plasma system treatment (flow rate: 15 sccm, time: 1 min, RF power: 20W) [Figure 2 (a-d)]. Alternatively, PDMS-coverslip bonding was also carried out using a thin PDMS (1:30 curing agent:PDMS) layer as the adhesive, followed by heating at 60^0^C for 4 hours [Figure 2 (e-g)].

**Figure 2:**
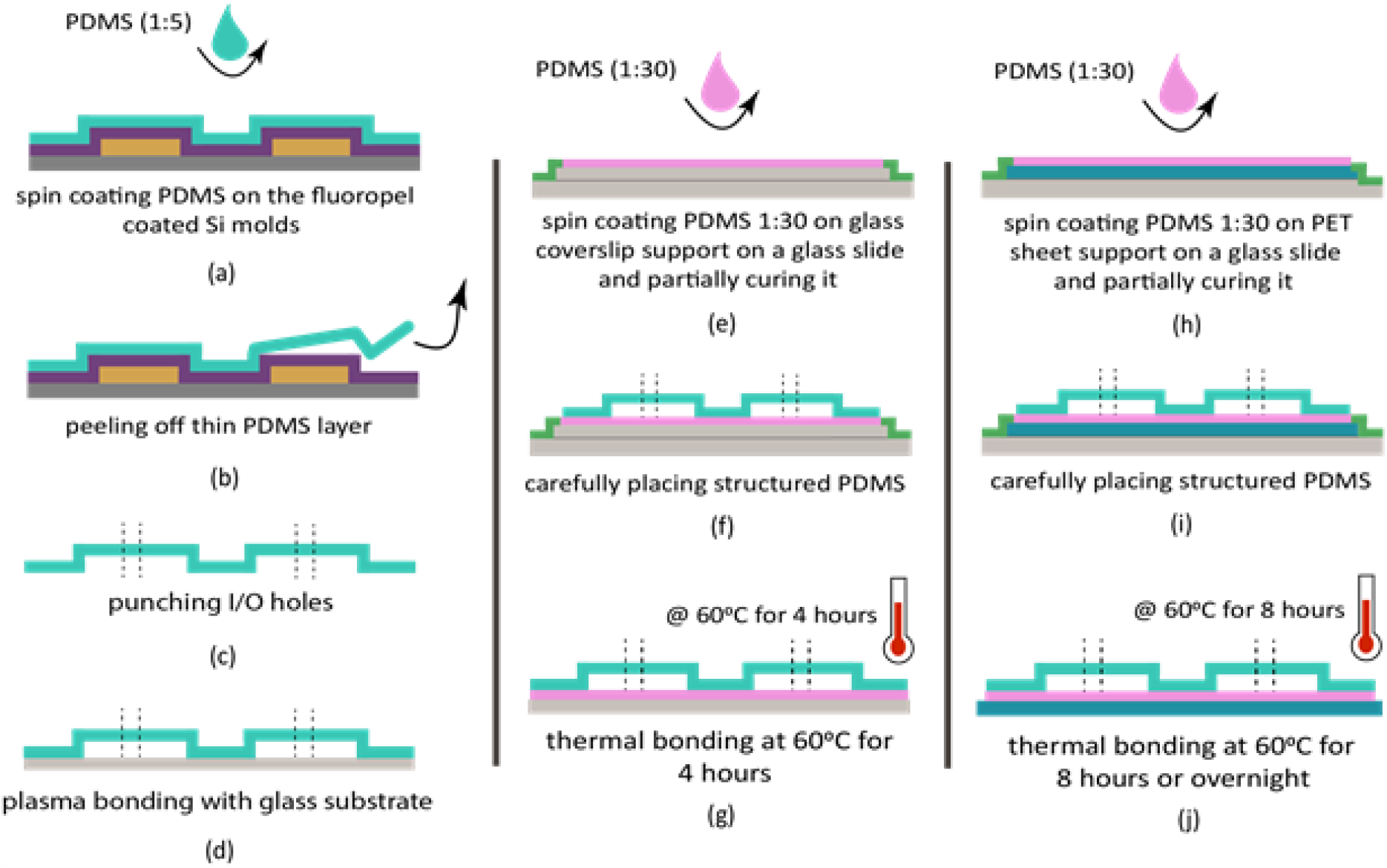
Fabrication process of 1-layer thin microfluidic device on glass coverslip (left and center) and flexible PET (right) substrates.

#### PET films as substrate

After degassing, PDMS (1:30) was spin-coated on the clean PET sheet at 6000 rpm for 60 s and was partially cured for 15 min at 60^0^C. The cured structured PDMS (1:5) was carefully placed on the partially cured PDMS (1:30) so as to prevent the clogging of the microchannels. But in this case, the heating was performed for 8 hours or overnight. The detailed recipe for 1-layer device fabrication is provided in the supplementary information.

### b) Fabrication of thin 3-layer device

For 3-layer devices, the flow and control channel molds were prepared on separate silicon wafers [Figure 3 (a-b)], and PDMS (1:5) was spin-coated at 500 rpm for 45 s [Figure 3 (c) left and center]. Once the thin PDMS layer on the FC and CC molds got cured, they were peeled off carefully [Figure 3 (d)]. For the membrane layer, PDMS was prepared in the ratio of 1:30 and degassed. This prepared PDMS was spin-coated on the fluoropel-coated bare silicon wafer at 6000 rpm for 3 min [Figure 3 (c) right]. After placing it on a flat surface for 5 min, it was heated at 80^0^C for 1.5 hours.

**Figure 3:**
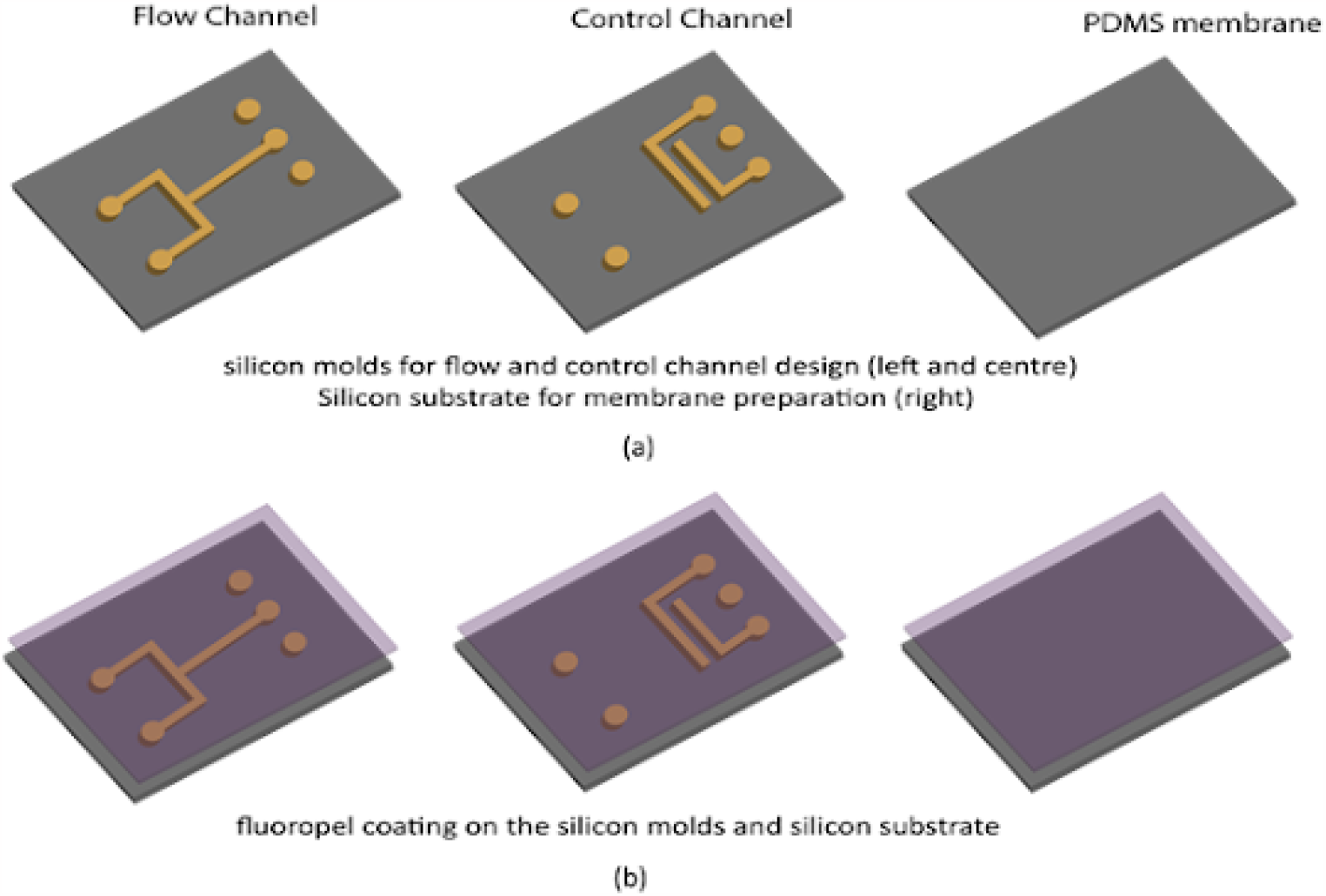
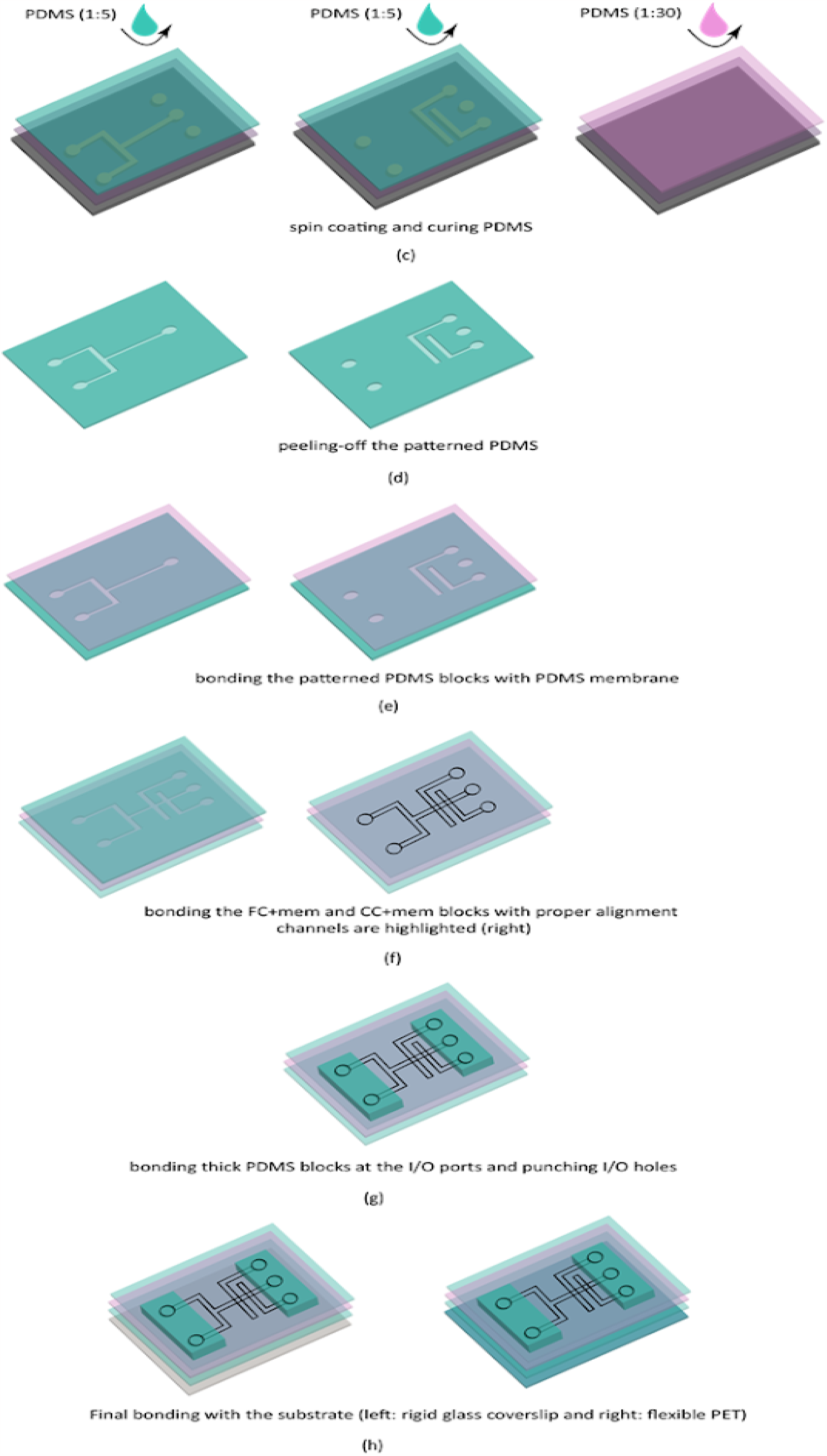
Fabrication process of 3-layer Quake-valve based thin microfluidic device on the glass coverslip and flexible PET substrates.

Now the thin membrane was bonded with FC and CC PDMS layers separately using oxygen plasma bonding such that the channels face the membrane [Figure 3 (e)]. We call these two units: FC+mem and CC+mem, respectively. This membrane prevents cross-contamination between the fluids in FC and CC. These units were bonded to each other using oxygen plasma treatment such that the membrane layers were sandwiched between FC and CC layers [Figure 3 (f)]. The alignment marks of both units were highlighted before bonding to ensure correct manual alignment. The layers were supported on a glass slide during bonding. After bonding, the glass slides were removed.

Hollow steel pins are commonly used to interface microchannels with tubing for fluidic flow. In conventionally used devices, the thick PDMS layer provides sufficient mechanical strength to fix the steel pin but the same does not hold true for the thin devices. Thus, we bonded thick PDMS blocks near the inlet-outlet ports to support the steel pins as in Figure 3(g). The final step involved providing mechanical support to the bonded PDMS layers. As shown in Figure 3(h), the glass coverslip and flexible PET sheet were used as the substrates, to fit a variety of applications.

### 2. Flow control setup

The main function of this controller is to monitor the pressure-driven flow into the microfluidics devices. A single-board computer (Raspberry Pi 4B) controlled the components of the setup. The compressed air (CA) flow in the CC was used to pressurize the membrane to function as a valve. A pressure regulator was connected to the input of a pneumatic valve terminal. The valve outputs were connected to CC inputs. A Python script executed on Rpi4 decided the selection and the duration of the actuated pneumatic valve. Another pressure regulator was connected to the fluid reservoirs to provide input to the FC. Alternatively, a syringe pump can be used to input the fluid into the FC at a controlled flow rate.

### Optical setup

The optical observation of the sample flow and regulation was performed with a zoom microscope (Eakins 2M camera microscope) at 4.5x magnification. The images were captured from the zoom microscope using Touplite software. In one set of experiments, Muscope was used to image the microfluidic channels. The sensor and light source in Muscope were controlled with the RPi 4B. All image processing was done offline on separate computers.

### 3. Sample preparation

#### a) L929 Mouse Fibroblast Cell Culture

Mouse fibroblast cells L929 (NCCS, Pune) were cultured in Dulbecco’s minimal essential medium (Gibco) supplemented with 10% FBS (Gibco) and 1% penicillin/streptomycin (Gibco) at 37°C in a humidified environment with 5% CO_2_. For monolayer culture, 5×10^4^ cells/cm^2^ were seeded in the culture dishes and routinely passaged upon 70% confluency.

For imaging, 5×10^4^ cells/cm^2^ cells were seeded on glass coverslips. Before cell seeding, coverslips were sterilized by autoclaving and pre-treated with ultraviolet light for 30 minutes. Upon 70% confluency, coverslips were processed for imaging experiments. Before imaging, cells were fixed with 4% paraformaldehyde (Sigma) for 15 minutes at room temperature. Then the cells were washed twice with 1X phosphate buffered saline (PBS) (Himedia) at room temperature and images were acquired under the phase contrast microscope under 20X and 40X objectives (EVOS™ M7000 imaging system).

#### b) Microbeads between two coverslips

The polystyrene microbeads of 10um diameter were thoroughly mixed in 1:10 PDMS. This mix was spin-coated on a clean glass coverslip supported on a glass slide. The coverslip is carefully taken off from the support glass slide and covered by another clean coverslip and heated at 80^0^C for 1.5 hours for PDMS to cure. This sample can be reused many times without any risk of any spill, debonding, or other contamination.

## Results and Discussions

The Quake valve can have 2-layer [24] and 3-layer [25] design architectures, based on the number of polymer layers used in their fabrication. The fabrication of a 2-layer microfluidics device involves spin-coating of PDMS on the photo-patterned molds. This hampers achieving the uniformly thick valves between FC and CC which causes unreliability in high-density chips as they involve multiplexing. On the other hand, in a 3-layer architecture, a thin PDMS membrane lies between the FC and CC layers. The deformation of the PDMS membrane controls the sample flow in the flow channel, upon the pressure exerted due to the air/liquid flow in the control channel. The thin membrane renders better thickness uniformity and therefore, enhanced density [25]. Moreover, it also offers to tune between membrane deformation *y* and applied pressure *P*, as given in eq. 1 for circular membranes [26]. Due to these merits, in this work, we chose to fabricate thin 3-layer microvalves.

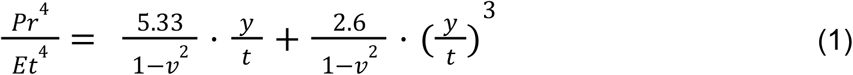

here, *E* is Young’s modulus and ν is Poisson’s ratio of the material used to make the circular membrane of thickness *t* and radius *r*.

Glass coverslips are the most common choice to fabricate thin devices. As the coverslip substrate is fragile to deal with, the flexible substrate is more appropriate for easy-to-handle and robust microfluidics devices fabrication, particularly for wearable bio-sensing applications. In this work, we have used the Polyethylene Terephthalate (PET) sheet as the flexible substrate [Figure 2 (h-j)].

Figure 5 shows the images of the active (3-layer) and passive (1-layer) microfluidic devices fabricated on rigid (glass coverslip) and flexible (PET) substrate using the discussed recipes. Figure 5 (I and III) shows the tilted, side, and top view of the thin and thick 1-layer and 3-layer devices on the glass coverslip (#1.5), respectively. On the other hand, top and side views of the thin and thick 1-layer and 3-layer devices on the flexible PET sheet are presented in Figure 5 (II and IV). The thickness of the thin and thick versions of both 1-layer and 3-layer devices differ by an order of 10s of micrometers. The thickness of PDMS layers was about 600*µ*m for the 3-layer device and can be further reduced.

We reported a simple procedure for PET-PDMS bonding without using any silane e.g. (amino propyl triethoxy silane, APTES or 3-Glycidoxypropyltriethoxysilane, GPTES), and/or chemical adhesives, and/or oxygen or ozone plasma treatments, unlike the reported recipes [27-30]. We used PDMS (1:30 curing agent:PDMS) as the adhesive layer followed by thermal bonding, in a similar way as that for the coverslips. The PET-PDMS bonding with our method performs equally well as glass-PDMS bonding using oxygen plasma treatment and was tested for pressure up to 2 bars. For the flexible valve devices, it is evident from Figure 5 (II) {c) and f)} and Figure 5 (IV) {c) and f)}, that the flexibility of thin devices is much more than that of thick devices. This makes thin devices an apt choice for flexible and wearable microfluidics applications.

Our process can be readily adapted to 2-layer devices, which are more common. As many researchers use 2-layer valve-based microfluidics circuits [3], we propose a simpler process to fabricate thin 2-layer push-down valves. This involves optimization of FC PDMS layer to form a thin membrane at the valve location. For e.g., in our case, where the channel height is about 15 *µ*m, the FC PDMS layer can be around 18*µ*m. The CC PDMS layer can be formed as mentioned in the previous discussions. These 2 layers can be appropriately aligned and bonded together with oxygen plasma or thermal bonding using PDMS as the adhesive.

During our efforts to use Muscope with microfluidics devices, we observed that the resolution of the DIHM depends on the quality of the captured holograms. The higher signal-to-noise ratio (SNR) of captured holograms yields better resolved real space reconstructed images. The SNR decreases as the thickness of the chip increases, consequently deteriorating the resolution of the Muscope. The Muscope’s magnification (M) and half-pitch resolution (Δx) are as given [31] –

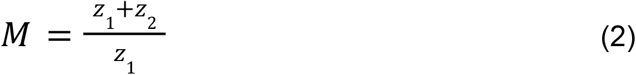

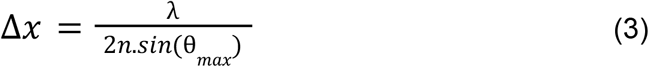

here, z_1_ and z_2_ are sample-source and sample-sensor distances, respectively. λ is the illumination wavelength, n is the refractive index and θ_max_ is the maximum scattering angle, such that

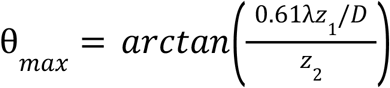

here, D is the lateral size of the light source.

Figure 6 presents the comparison between the holograms of thick and thin 1-layer devices on the coverslip substrate. The holograms captured with Muscope, for the two devices, are presented in different working conditions namely when the channels were empty [Figure 6(a and e)], PBS filled in the channel using a syringe pump with a flow rate of 10um/min [Figure 6(b and f)], L929 cells present in the channel [Figure 6(c and g)]. The holograms of L929 cells are present inside the red dotted boxes in Figure 6(c and g). For thick microfluidic devices, the recorded holograms were not rich enough to result in a high-resolution image of the sample. The fringes corresponding to the higher frequency components were either faint or totally absent. The holographic reconstruction algorithm [31] was used to retrieve amplitude and phase information from the recorded holograms for z_2_=350*µ*m. We validate the necessity of thin microfluidics devices by comparing the reconstructed amplitude image of the portion of holograms enclosed in the yellow box. The reconstruction in the case of the thick device is very dim and the presence of the cells is very unclear [Figure 6(d)]. On the other hand, in Figure 6(h), for the thin device, the reconstruction of the channel and cell is much better.

A similar trend in the hologram quality was observed for the flexible 1-layer devices on the PET substrate (Figure 7). The high-quality hologram, with more and better contrast, was recorded for the thin version of the 1-layer devices as in Figure 7 (a and c). The reconstructed images of the 50*µ*m wide channel filled with PBS for the two devices are highlighted within the yellow lines in Figure 7 (b and d) for z_2_=300*µ*m. For the thick device, the channel is hardly recognizable, whereas is clear for the thin device. Also, notice that the decreased source-sample distance in thin devices resulted in a slight enhancement in the magnification (eq.2). The presence of circular fringes outside the microchannel in Figure 7 is due to the presence of small dots on the PET substrate used in this work. Hence, it is recommended to use a good-quality PET sheet to fabricate the devices.

The retrieval of the complete sample field (i.e. amplitude and phase) using holographic reconstruction was used to extend the Muscope’s capability to function as a PCM with no modification in the hardware. The obtained field was digitally modified following the pseudocode [32]

- *determine the zero frequency component of the complex sample field*
- *apply* π*/2 phase shift to it*
- *find the intensity of this mutated complex field*

The Muscope was used to image L929 cells fixed between two coverslips. The captured hologram and its reconstructed phase information are presented in Figure 8 (a and b). In Figure 8 (b and c), an improvement in the image contrast can be clearly visualized. It was also observed that the digitally retrieved phase contrasted image highly depends on the efficiency of the reconstructed field. A more efficient and/or super-resolution technique can be utilized to enhance the accuracy of the retrieved phase information. On comparing the digital phase contrast image with that obtained using conventionally used PCM (20x magnification), we affirm the Muscope’s potential as a phase contrast microscope without imposing any additional hardware accessories compulsions. We also presented Muscope-based PCM for the L929 cells in the 50*µ*m wide channel of 1-layer microfluidic device in Figure 9. The contrast enhancement for the digital phase contrasted image using Muscope is much better than the reconstructed phase image.

The controller set-up shown in Figure 4 was used to test the working of the thin 3-layer microfluidics devices fabricated with our process. The microvalve functioning, in both rigid and flexible devices, was affirmed by the optical observation using the zoom microscope (Eakins 2M camera microscope) at 4.5x magnification in Figure 10 and 11. Figure 10 (a) and Figure 11 (a) show the valve in OFF state whereas Figure 10 (d) and Figure 11 (d) show the actuation of the valve i.e. ON state. The valves were actuated using compressed air (CA) in the CC. The valve appears strained and relaxed in the presence and absence of CA, respectively. The valves were tested to regulate IPA and microparticles flow in the channels. In Figure 10 (b), IPA can be seen to flow upward towards the valve but once the valve was turned ON, as in Figure 10 (e), IPA flow experiences an increased resistance and thus, its flow was impeded. The same holds true in Figure 10 (c and f) for the flow of microparticles. A similar approach was used to test fluidic regulation in the flexible devices, as shown in Figure 11 (b and e) and Figure 11 (c and f) for IPA and microparticles, respectively. All the devices were tested for pressure up to 1.8 to 2 bars even after 5-6 months of fabrication without any degradation in the device operation. This affirms that our process flow can be employed to fabricate the thin devices for active and passive microfluidics devices.

**Figure 4:**
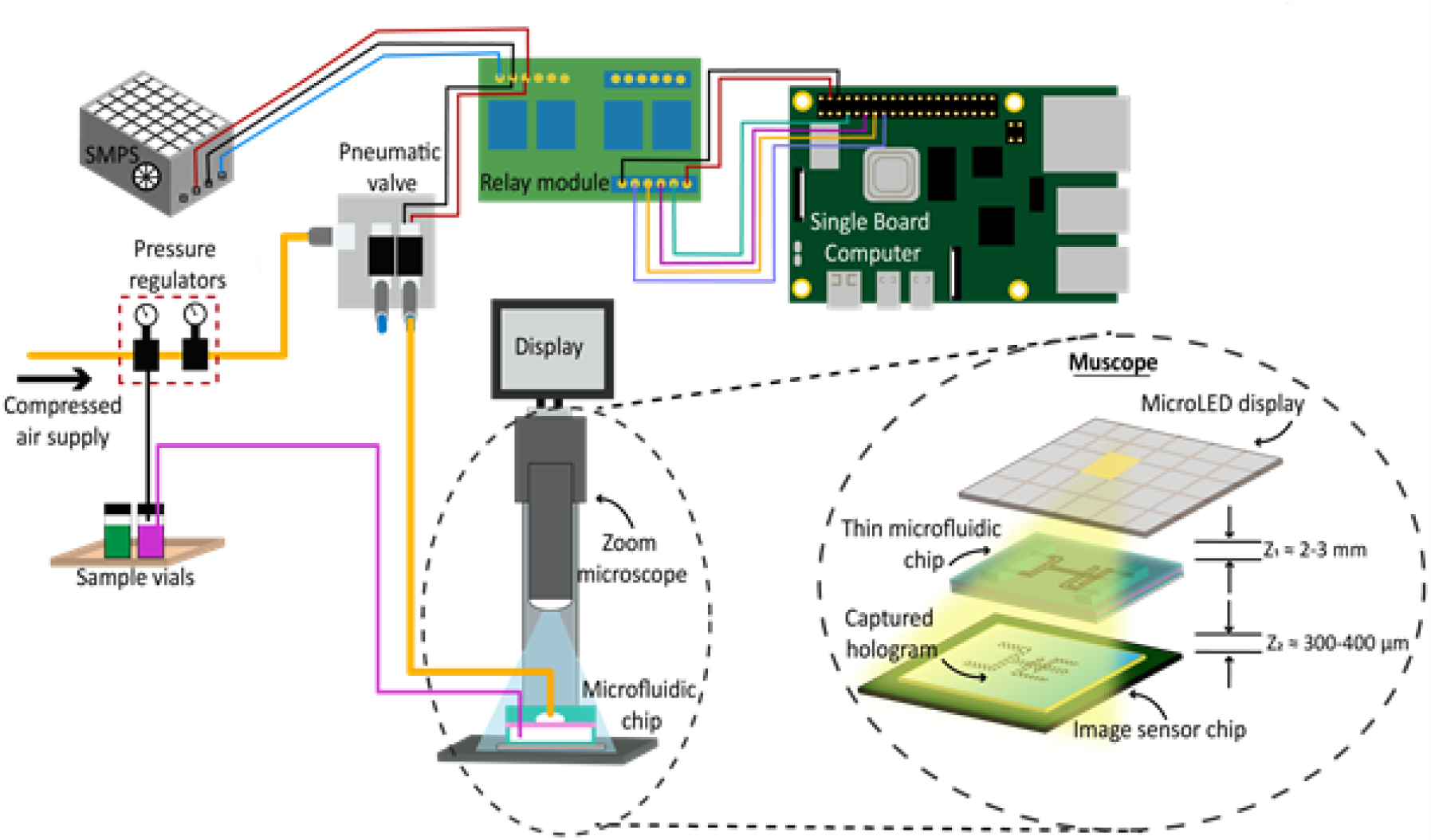
The microfluidic controller setup. The table-top zoom microscope is used for optical observations. It can be replaced by Muscope, resulting in system miniaturization.

**Figure 5:**
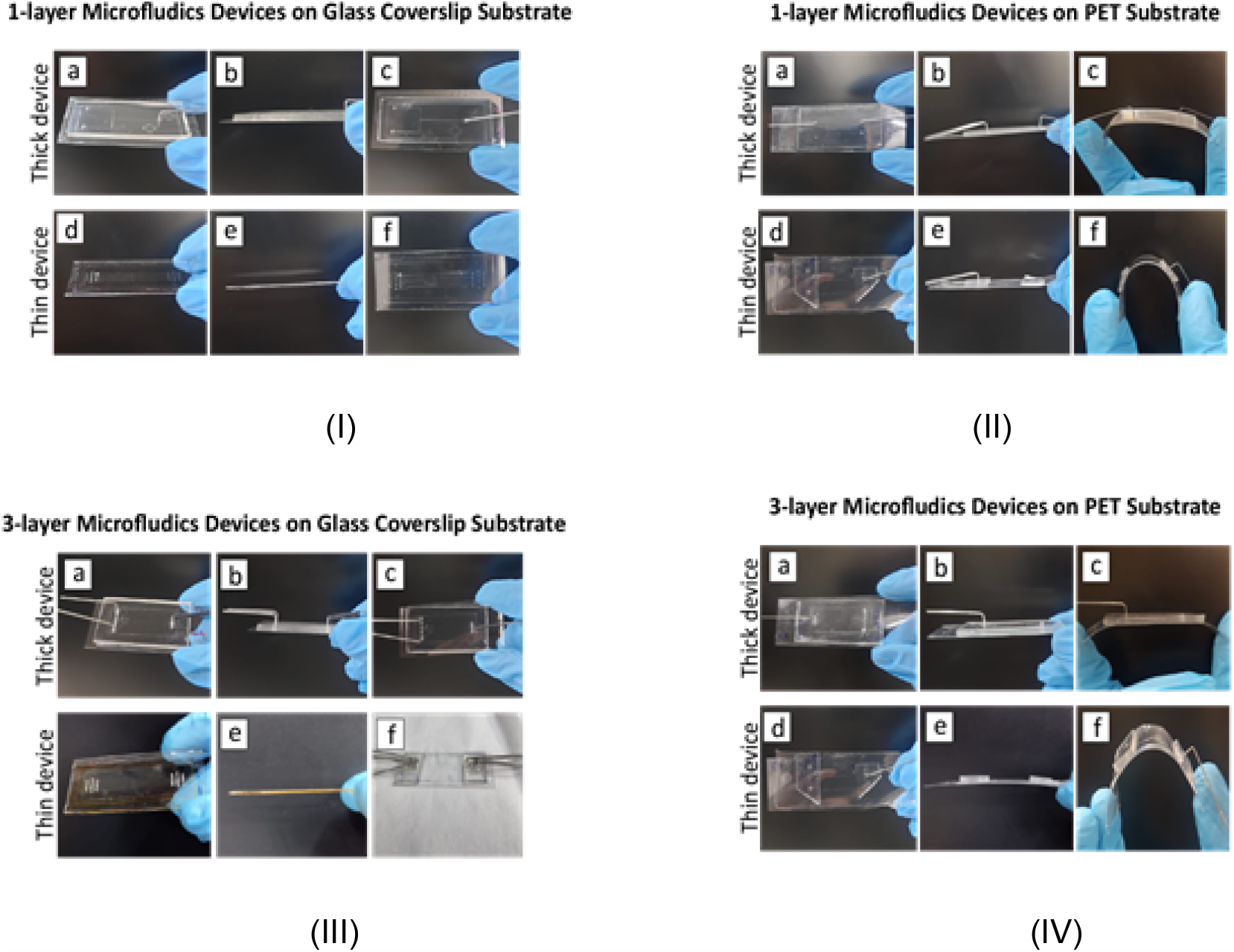
Device images to compare the thin and thick devices. The images on the left show the 1-layer and 3-layer devices on the rigid coverslip substrate. The images on the right show the 1-layer and 3-layer devices on the flexible PET substrate.

**Figure 6:**
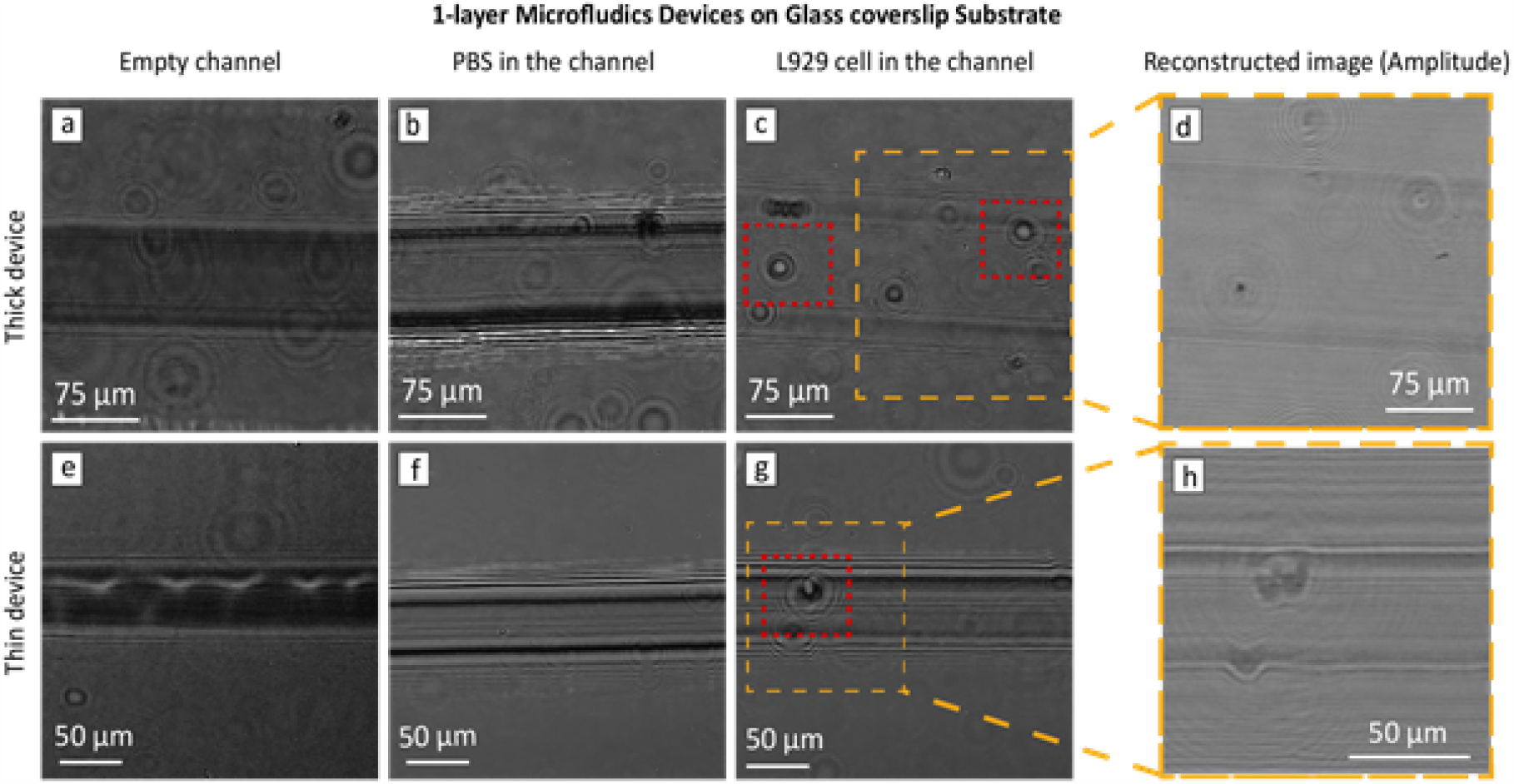
Comparison of thick and thin 1-layer device images, captured with Muscope in different working conditions. (a) and (e) present the channels in the empty state, (b) and (f) present when the PBS was filled in the channels. The L929 cells were made to flow in the channels shown in (c) and (g). The presence of L929 cells is indicated by the red box. The comparison of the holographic reconstructed images of the portion of the images in (c) and (g) for z_2_=350*µ*m is presented in (d) and (h).

**Figure 7:**
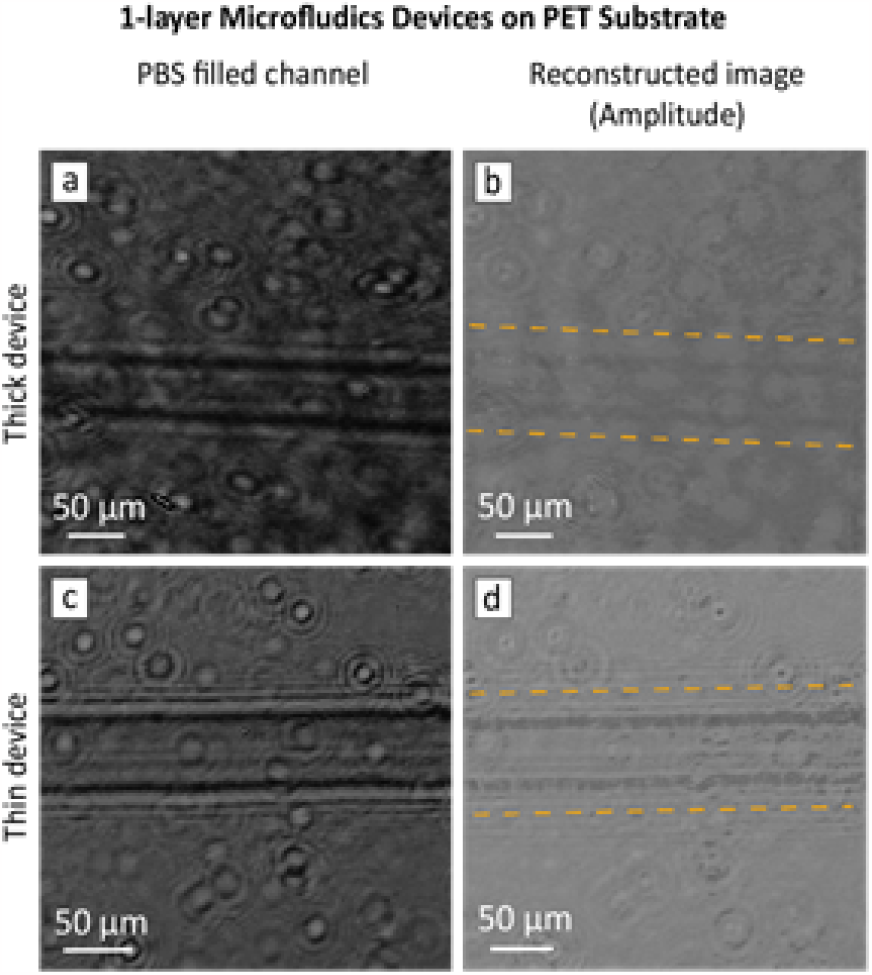
The comparison of holograms capture when using Muscope to image 1-layer thick and thin devices on PET substrate.

**Figure 8:**
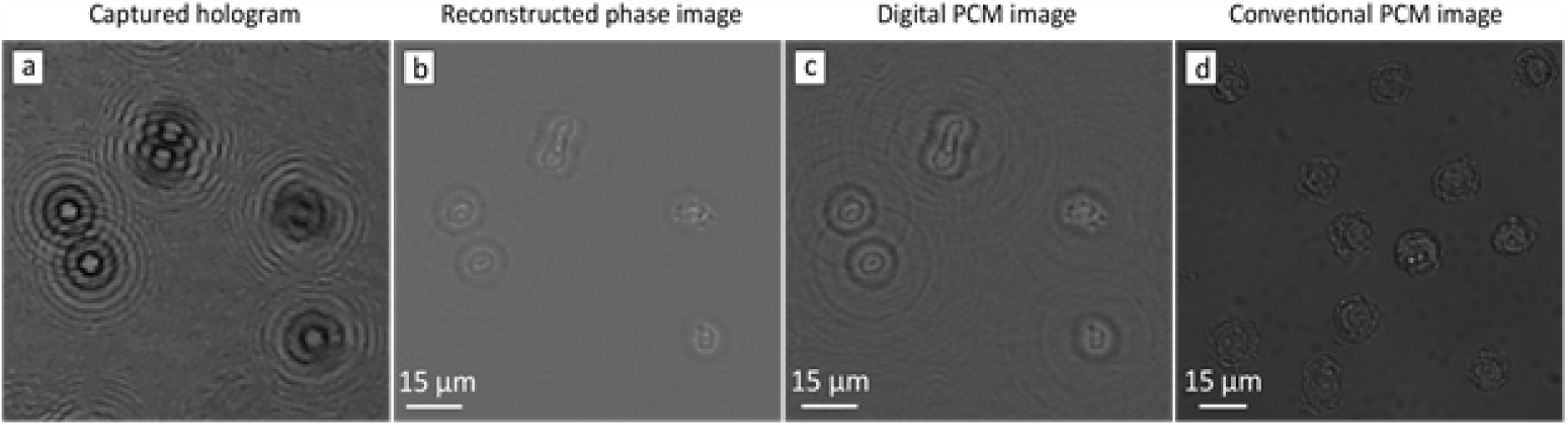
presents the Muscope’s function as PCM. (a) shows the hologram of L929 cells, captured with Muscope, (b) shows the phase image obtained using holographic reconstruction, (c) PCM image of L929 cells by digitally mutating reconstructed sample field, and (d) the L929 cells image obtained with conventional PCM at 20x magnification.

**Figure 9:**
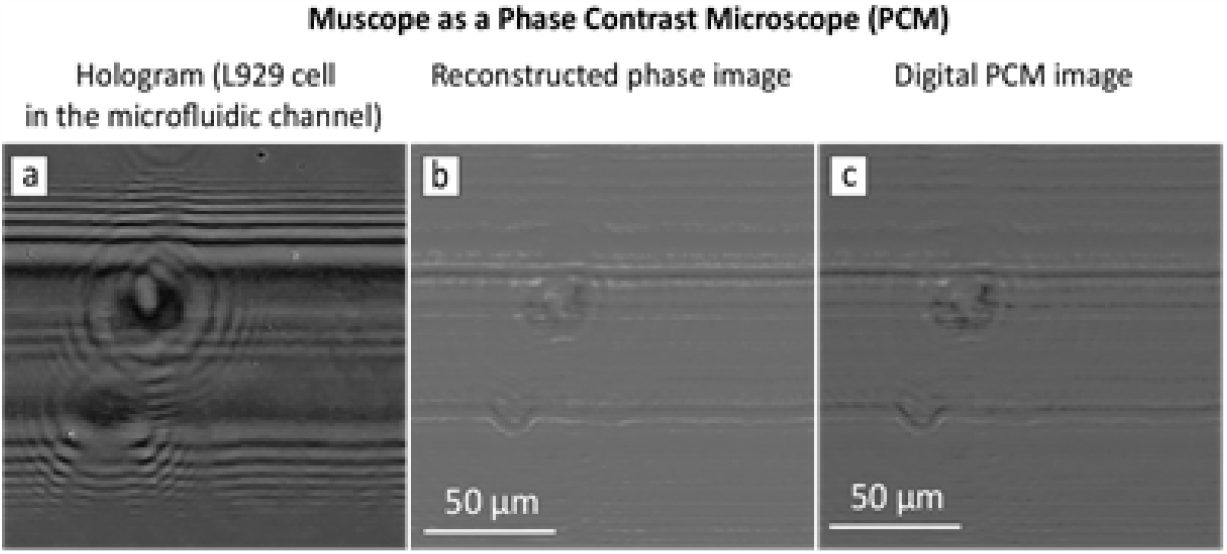
presents the Muscope’s function as PCM. (a) shows the hologram of L929 cells in the 50*µ*m wide microfluidic channel, captured with Muscope, (b) shows the phase image obtained using holographic reconstruction, (c) PCM image obtained by digitally mutating reconstructed sample field.

**Figure 10:**
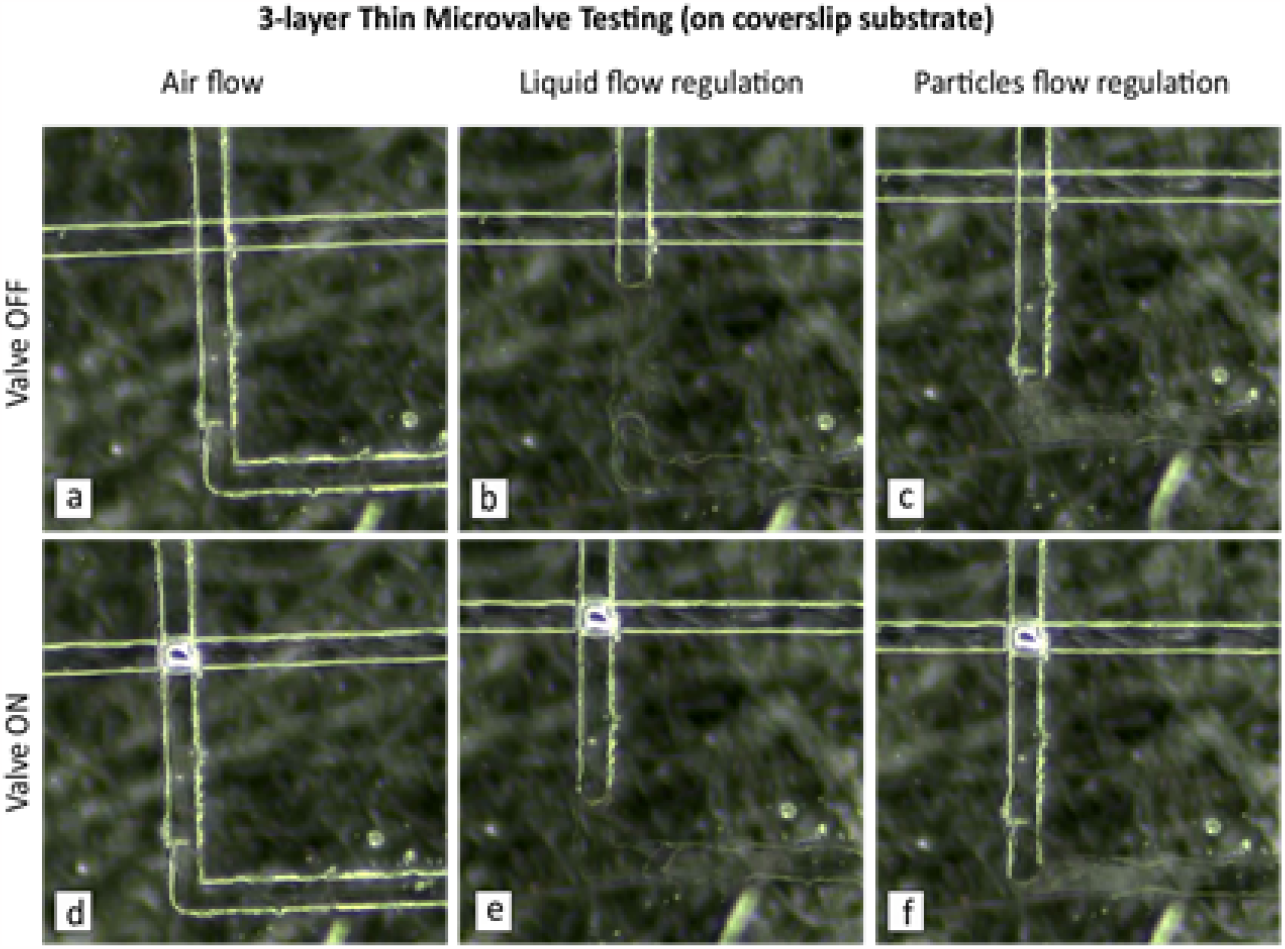
Testing valve action for the thin 3-layer device on coverslip substrate. (a) and (d) present the valve in the absence and presence of CA in the CC, respectively. (b and e) and (c and f) show the IPA and microparticles flow regulation, respectively.

**Figure 11:**
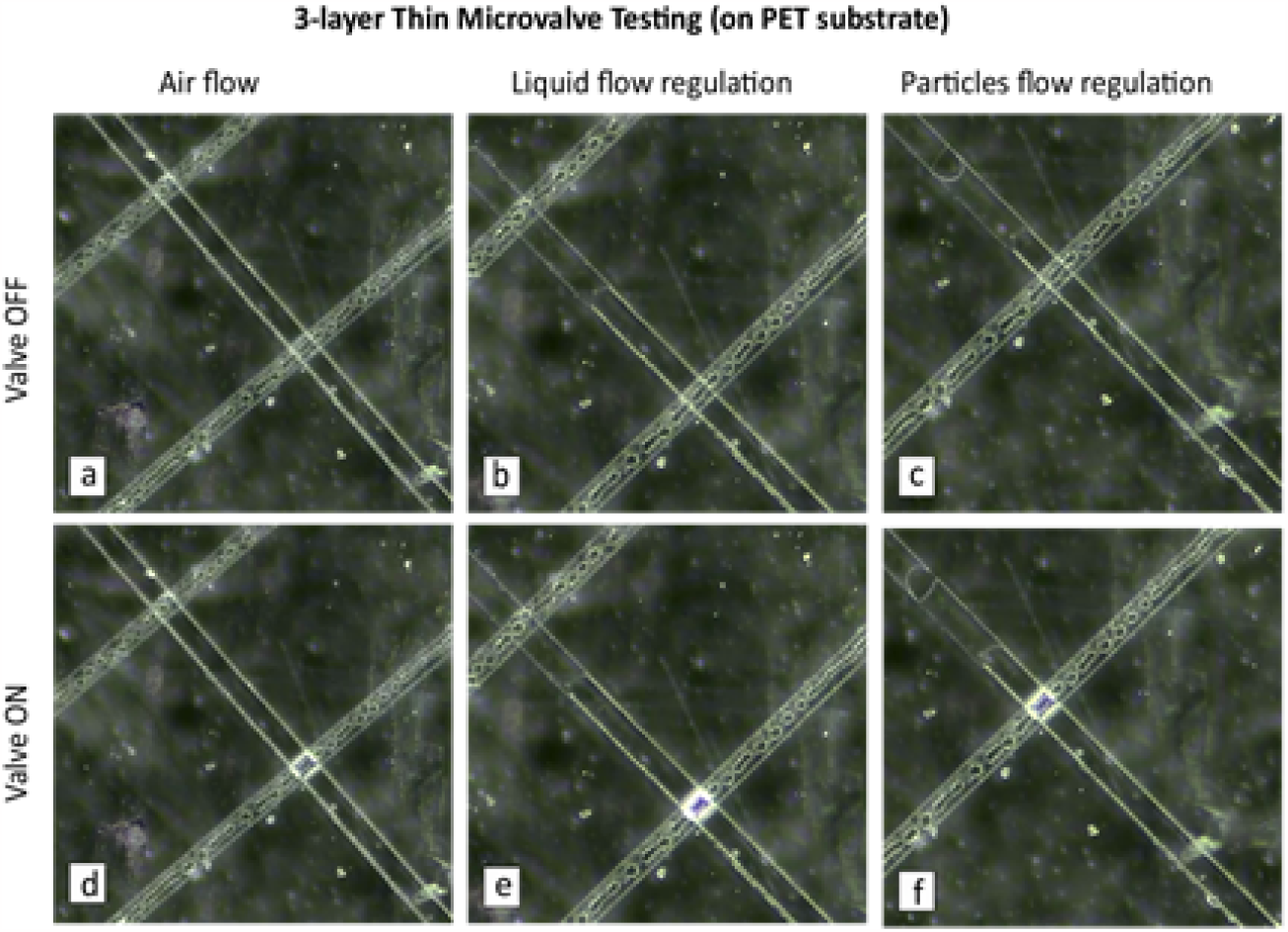
Testing valve action for the thin 3-layer device on PET substrate. (a) and (d) present the valve in the absence and presence of CA in the CC, respectively. (b and e) and (c and f) show the IPA and microparticles flow regulation, respectively.

To experiment with multiple valves, we tested the working of two valves by actuating them simultaneously. Figure 12 (a) and (b) show the two valves in OFF and ON state, respectively. When the two valves are actuated together, better fluidic flow regulation is observed. This concept can be extrapolated to achieve flow cytometry. Also, by having 3 valves together and actuating them in an appropriate sequence, they can act as a micropump. Thus our proposed fabrication methods are suitable for large scale microfluidic circuit chips for high throughput biology.

**Figure 12:**
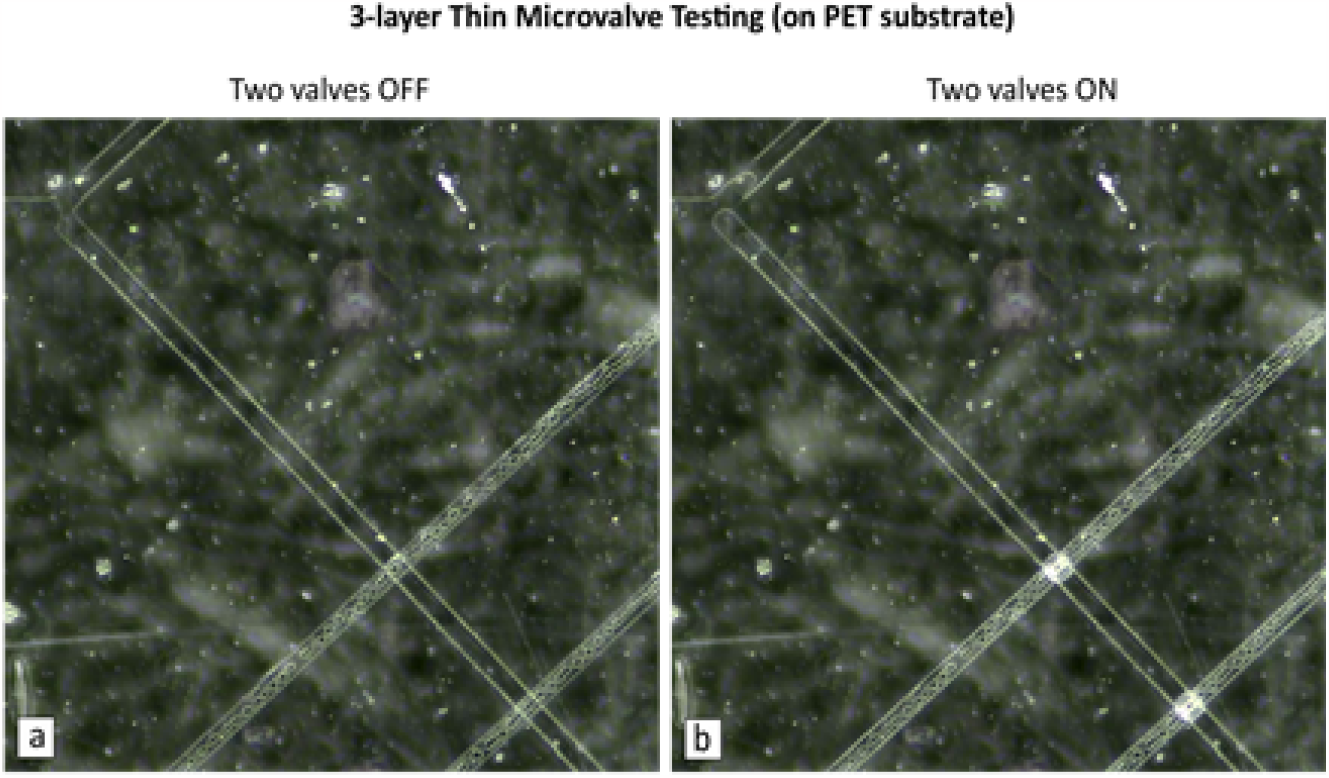
Testing liquid flow control using multiple valves in the thin 3-layer device on PET substrate.

## Conclusion

We reported the novel soft-lithography based processes to fabricate thin active and passive microfluidic devices for sample space-limited applications. One such application arises by the replacement of traditional lens-based microscopes, used in mLSI, with their compact lensfree equivalents. We demonstrated the requirement for thin microfluidic devices with our recently reported lensless DIHM, termed Muscope. The employment of Muscope with mLSI can lead to the development of a new generation of compact microfluidic integration systems. We also fabricated flexible single- and multi-layer microfluidic devices on PET substrate. We confirmed a simple method for PDMS-PET bonding without the use of any silane group, chemical adhesive, or plasma treatment. This opens a huge scope to develop convenient-to-handle, robust, and compact wearable microfluidic-based sensing technologies like biosensors, pressure sensors, etc.

The 3-layer Quake valve has many merits over its 2-layer counterpart, thus, was our choice for this work. We also affirmed the successful functioning of 3-layer thin device on the rigid and flexible substrate to govern the flow of liquid (IPA) and microparticle suspension. Our devices had rectangular FC cross-sections, which can be made semi-circular using the reflow technique during mold preparation, ensuring perfect valve switching. All our devices were tested for pressure up to 2 bars even after 6 months of fabrication. The breakdown pressure and shelf-life of the devices can be tested using the peel test, blister test, etc. Also, we proposed a simpler process flow to fabricate the thin 2-layer Quake valve. The successful working of multiple valves confirms our fabrication process to be adapted for complex mLSI circuit chips.

We also demonstrated the potential of Muscope as a PCM for samples on the coverslip and inside the microfluidic channel. The more accurate sample field retrieval leads to the better phase contrasted image obtained with Muscope. This is possible to achieve by using more efficient holographic reconstruction and/or super-resolution techniques.

## Acknowledgment

We are grateful to the Science and Engineering Research Board (SERB), DST, Government of India, for providing financial aid for this research work. We are thankful to our labmates Ms. A.K. Niketa and Ms. Saima Hamid for their support. We would also like to extend our sincere gratitude to Dr. Satish Bonam, Mr. Nitish Kumar Singh, and Mr. Naveen Kumar Kamatchi Ramakrishnan for their help in performing the experiments.

