## Supplementary Information for "Thin Microfluidic Chips with Active Valves"

[a] Department of Electrical Engineering, Indian Institute of Technology Hyderabad, India
[b] Department of Biomedical Engineering, Indian Institute of Technology Hyderabad, India

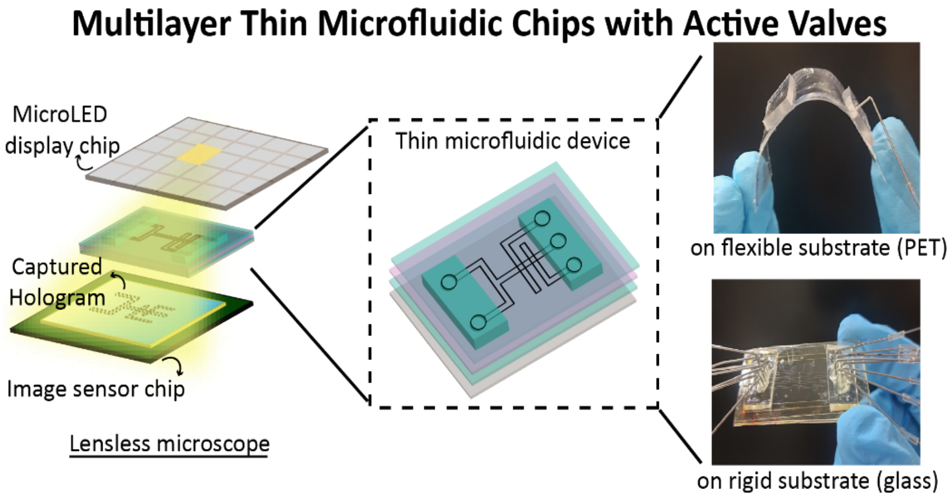

### Contents

I. Fabrication of silicon molds
II. Fabrication of thin 1-layer devices

---

#### I. Fabrication of silicon molds

The microchannel design for each PDMS layer of the devices was transferred to separate silicon wafers using standard photolithography. This process included: a) chrome mask preparation b) pattern transfer to the silicon substrate c) preventing delamination of the patterned photoresist from the silicon substrate in the further steps.

Firstly, the desired microchannel patterns were designed using the gdspy library of the python scripting language. Alternatively, the other software such as KLayout, Clewin, LayoutEditor, LASI, AutoCAD etc. can also be used. The cleaned chromium coated glass plates were used as the substrate to prepare the photomasks. The novolak based positive photoresist S1813 was spin-coated on the cleaned glass plate to result a 1-2 micron thick layer. It is then soft baked for a minute at 120$^0$C. Then the mask was written using the laser writer machine LW 405B. Once the pattern writing was over, the photoresist was developed using the developer MF319 for about a minute and then immediately cleaned with DI water. Once the glass plate is dehydrated well, the chromium etching is performed and the patterns are checked under the optical microscope. Once the desired patterns are obtained, the residual S1813 was removed with acetone and IPA wash. The masks acted as a blueprints of our design.

The second step was to prepare master/mold using conventional photolithography. The epoxy-based negative photoresist (NPR) SU8 2015 was spin-coated on the clean silicon substrate to result 13-15 um thick layer. An optional step prior to spin coating is to treat silicon wafer with $O_2$ plasma to promote its adhesion with SU8 NPR. Then prebaking was performed at 95$^0$C to remove the solvent. The wafer was then exposed to 365 nm UV radiation using the mask aligner. The datasheet was followed to finalize energy dosage, post exposure bake and development of the SU8 2015. It is advisable to check for the desired patterns under the optical microscope after development. Finally, once the desired microchannel patterns were achieved, the sample was hard baked for 20 min @150$^0$C.

Thirdly, to avoid the delamination of SU8 2015 in further steps (in particular, replica molding), we used Fluoropel 800 0.2%. Fluoropel was spin-coated on the molds at 3000 rpm for 30 sec to result a hydrophobic and oleophobic layer on the molds' surfaces. The molds were then left to air dry for 5 min, followed by heating at 120$^0$C for 20 min. Fluoropel coating is a safer substitute for the conventionally used Tetra Methyl Chloro Silane (TMCS) for mold silanization. Unlike TMCS, fluoropel spin coating is performed a single time and the molds can be used for replica molding for many months, ensuring an easy release of very thin layer of PDMS. This avoids the requirement to silanize the molds with TMCS each time before performing the replica molding.

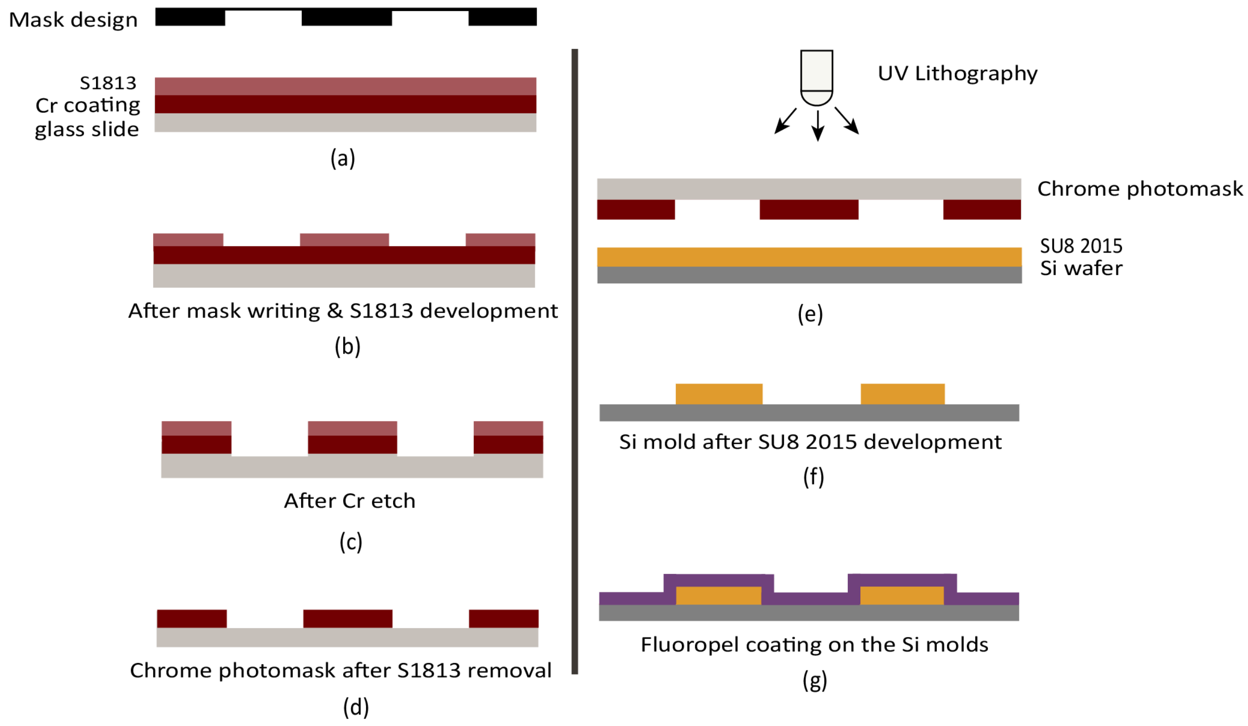


Figure 1: The process flow for chrome mask [(a)-(d)] and silicon mold preparations [(e)-(g)].

#### II. Fabrication of thin 1-layer devices

We spin coated the PDMS (1:5) on the silicon molds and cured it at $80^0$C for 1.5 hours. The thin PDMS layer was carefully peeled off the molds. Care should be taken to prevent tearing of thin PDMS layer. The inlet-outlet ports were pierced with the puncher. In this work, the structured PDMS was bonded with the glass coverslip (#1.5) using the oxygen plasma system.

Another simpler method to bond PDMS with the coverslip is to use a thin PDMS layer as the adhesive. To demonstrate this, a thin layer of PDMS (1:30 curing agent:PDMS) was spin coated on the coverslip at 6000 rpm for 60 sec and was partially cured for 15 min at $60^0$C. The structured PDMS was carefully placed on the partially cured PDMS so as to prevent the clogging of the microchannels and was left for heating for 4 hours at $60^0$C to thermally bond with the glass coverslip. The two bonding methods worked well and were tested for pressure up to 2 bars.

As coverslip is fragile to handle, a flexible substrate can act as a suitable substitute for easy handling and also for wearable electronics. In this work, we have used the Polyethylene Terephthalate (PET) sheet as the flexible substrate. The method opted to bond the structured PDMS with PET is similar to that of the coverslips. We used 1:30 curing agent:PDMS as the adhesive layer in the similar way as that for the coverslips. But in this case the heating was performed for 8 hours or overnight.

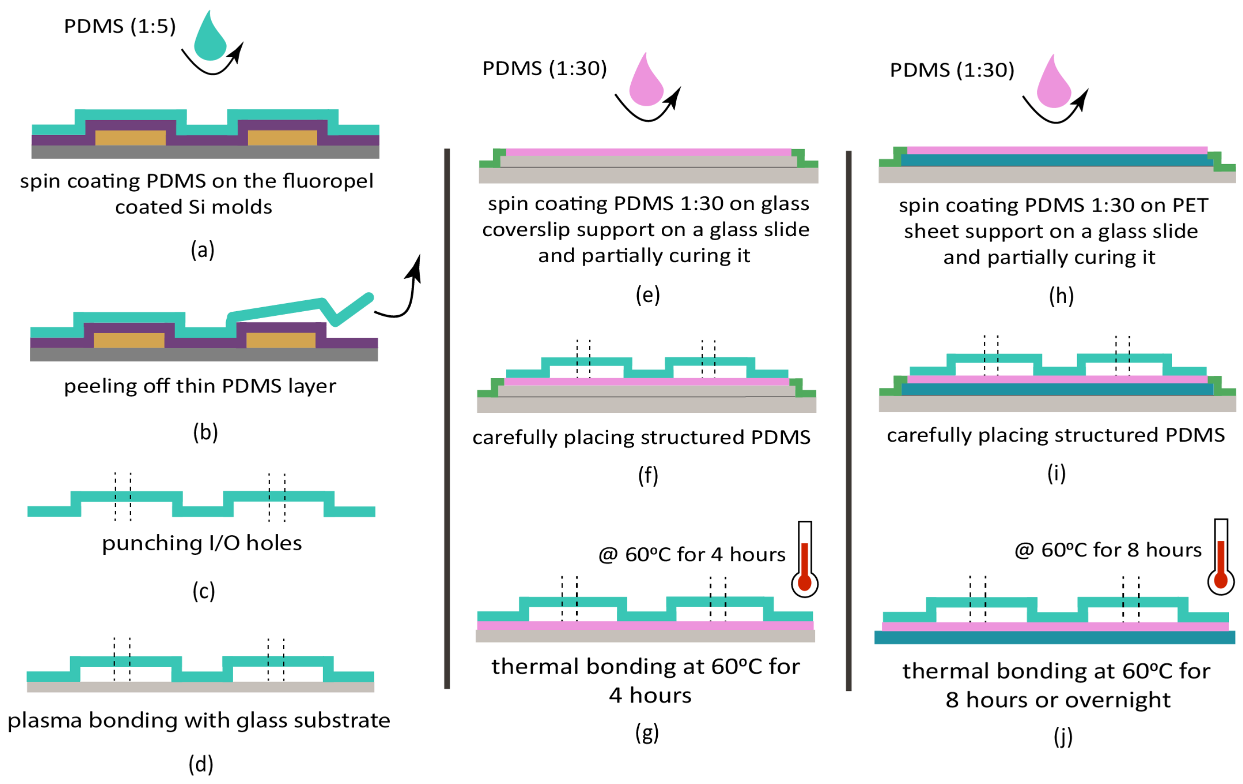


Figure 2: Fabrication process of 1-layer thin microfluidic device on glass coverslip (left and center) and flexible PET (right) substrates.
